# Real-time affinity measurements of proteins synthesized in cell-free lysate using fluorescence correlation spectroscopy

**DOI:** 10.1101/2024.09.28.615564

**Authors:** Chao Liu, Steven A. Hoang-Phou, Congwang Ye, Emma J. Laurence, Matthew J. Laurence, Erika J. Fong, Nikki M. Hammond, B. Dillon Vannest, Nicholas N. Watkins, Ted A. Laurence, Matthew A. Coleman

## Abstract

Rapid, high throughput measurements of biomolecular interactions are essential across medicine and bioscience. Traditional methods for affinity-screening proteins require a long and costly process involving cell-based expression, purification, and titration of multiple concentrations to arrive at a binding curve. In contrast, we have developed a fast and simple approach that yields a wealth of information about the expression of the protein and its binding characteristics, all in a “one-pot reaction” and done in under several hours without the need for protein purification. The method uses cell-free protein synthesis to produce the protein of interest in the presence of its binding partner, while simultaneously using fluorescence correlation spectroscopy (FCS) to measure the increasing concentration of the protein and its binding to the binding partner. We characterize the sensitivity limits of this method by measuring the binding between the green fluorescent protein (GFP) and a low picomolar-affinity anti-GFP antibody and found that we can quantify *K*_D_s down to the high picomolar to low-nanomolar range. We further demonstrate the method in a potentially ultrahigh-throughput sample format, in which FCS measurements are collected inside microcapsules. This work lays the foundation for a platform aimed at production and in situ affinity screening of thousands of different proteins.

## INTRODUCTION

High throughput screening is critical for developing therapeutics for new and reemerging threats from pathogens ^1^. Strategies for increasing the efficiency and speed of this screening are valuable for testing candidate therapeutics, which may include libraries of combinatorial/random compounds and computational designs ^2,3^. In this work, we develop a strategy for reducing the steps in the screening process. Conventionally, the production of proteins is separate from a screening step measuring interactions between proteins. Here we introduce a strategy in which we measure binding affinity for protein-protein interactions simultaneously with the production of the proteins using fluorescence correlation spectroscopy (FCS). Importantly we combine FCS with coupled transcription and translation cell-free protein synthesis (“cell-free” for short) lysates, which enable the production of proteins encoded by plasmid DNA *in vitro*. We can assess the concentration of produced protein as it increases while also measuring binding to its binding partner in solution. With this strategy, proteins can be designed, synthesized, and screened efficiently all in “one-pot reactions”, in which the entire process for each protein is completed in a single sample solution.

FCS is sensitive to individual fluorescently labeled molecules diffusing into and out of an open detection volume defined by tightly focused laser beams and a confocal pinhole ^4^ ^5^ ^6^. There are two basic ways to measure protein interactions with FCS. In a single-color experiment, an increase in diffusion time related to the increase in size of the diffusing molecule due to binding is used to measure binding. In a dual-color experiment, binding is measured using cross correlations that only occur when proteins with two different colored labels are bound to each other ^7^. While the strategy presented here potentially could be used in both single-color and dual-color experiments, here we focused on single-color experiments. The benefits of FCS for these measurements include sensitivity to low concentrations while being able to perform the measurements in a solution environment without washing or purification steps.

We use *E. coli* cell-free lysates to rapidly synthesize proteins directly in our sample wells during FCS observation, bypassing the need for laborious cell-line generation and protein purification ^8^ (Figure 1). Previously, it was shown that FCS can be performed on proteins produced in cell-free lysates ^9^ ^10^ ^11^. Here, we also demonstrate the ability to obtain a full binding curve by tracking fluorescent protein expression directly within the cell-free reaction. By combining cell-free protein synthesis and FCS, we simultaneously perform the synthesis and affinity measurement steps, thus enabling a simpler, faster workflow for screening applications.

**Figure 1.**
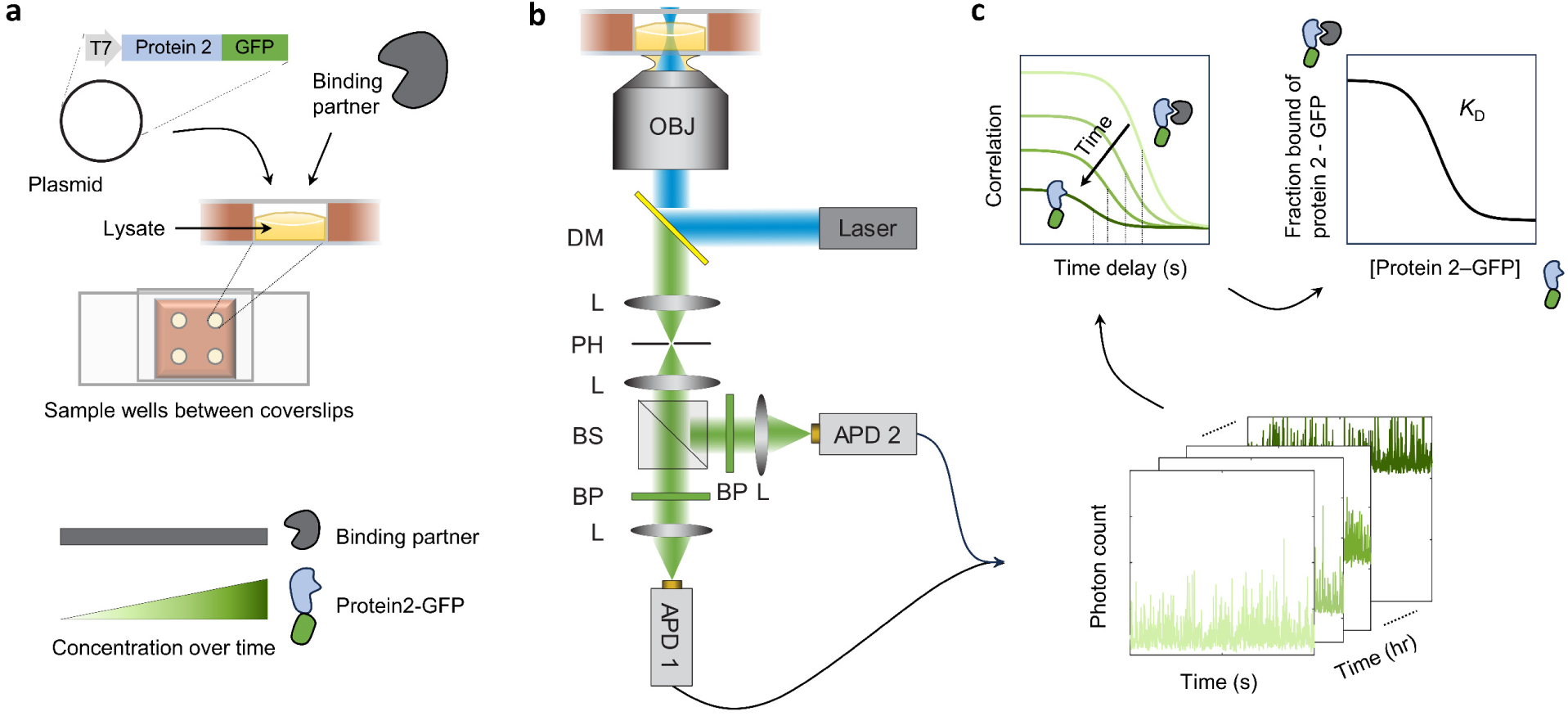
Experimental setup to measure binding of protein during cell-free synthesis by fluorescence correlation spectroscopy (FCS). **a.** Reactions consist of the cell-free expression lysate, the protein of interest (“Binding partner”), and a plasmid encoding the fluorescently tagged second protein (“Protein2-GFP”) that is expected to bind to the Binding partner. Reaction solutions (∼ 6 µl) are incubated in sample wells between coverslips, keeping a layer of air above. Different sample wells may contain different variants of Protein2-GFP when screening Protein2 for binding, for example. The concentration of Protein2-GFP increases over time as the expression proceeds while that of the Binding partner remains constant. **b.** Samples are measured over time by simple one-color FCS. A 488nm laser excites the GFP, and emitted photons are selected by a confocal pinhole (PH) and detected by two avalanche photodiodes (APDs). OJB: objective, DM: dichroic mirror, L: lens, BS: beamsplitter, BP: bandpass filter. c. Bottom: data consists of arrival times of each photon on the detectors, collected for different samples over several hours of protein synthesis. Top left: calculated correlations are expected to shift down and left over experimental time, as amplitude decreases due to increasing protein concentration while overall diffusion time decreases due to an excess of free Protein2-GFP over its Binding partner. Top right: a two-species fit to correlations yields the fraction of Protein2-GFP bound and its concentration, producing a binding curve from which the binding affinity (*K*_D_) is determined.

### System overview

Our method uses FCS to measure production and binding of fluorescently labeled proteins in cell-free lysates in a one-pot reaction without the need for protein purification (Figure 1). Each reaction required only ∼6 µl consisting of the cell-free lysate, one purified protein of interest (“Binding partner”), and a plasmid encoding a fluorescently tagged (e.g., green fluorescent protein, GFP) second protein of interest (“Protein2-GFP”) (Figure 1a). Each reaction solution is incubated at room temperature as a thin layer in a sample well between two coverslips, keeping a layer of air above. The layer of air ensures spatially uniform production and detection of the GFP signal across the well over time since oxygen is required for both protein synthesis ^12^ ^13^ and GFP maturation ^11^ ^14^. Indeed, sample chamber configurations in which only a portion of the solution was in contact with air exhibited a spatial gradient of GFP signal (Supplementary Figure 1). Over the course of protein synthesis, the concentration of Protein2-GFP increases while that of the Binding partner remains constant, so a series of measurements over time produces different concentrations as required for a binding curve.

The fluorescent signal in each sample well is measured over time by one-color FCS (Figure 1b). A 488nm laser excites the GFP, and emitted photons are selected by a confocal pinhole and detected by two avalanche photodiodes. Over several hours of protein synthesis, an automated stage cycles through each sample well to collect 30-second segments of data consisting of photon arrival times at each detector (Figure 1c, bottom). The *g*^(2)^ correlation is calculated as the cross-correlation of the two photon streams to mitigate the nano-to micro-second scale peak due to detector afterpulsing ^15^. As Protein2-GFP is generated over time, the correlation amplitude decreases, and the concentration can be determined from the amplitude given the calibrated focal volume (Figure 1c, top). Importantly, the overall diffusion time is initially high since the small amount of Protein2-GFP would be bound to its Binding partner but decreases over time as free Protein2-GFP is produced in excess of the Binding partner. A two-species fit to correlations yields the fraction of Protein2-GFP bound, which, together with its measured concentration, produces a binding curve from which the binding affinity (*K*_D_) is determined.

To demonstrate the feasibility of our method and to investigate its requirements and limitations, we studied the binding of GFP to a commercially available anti-GFP antibody. As antibodies can have very high affinities (low *K_D_*s), this choice of model proteins enabled us to determine the sensitivity or lower detection limit of the method. We first show that FCS can measure the fluorescent signal in the background of cell-free lysate and thus track the production of GFP over time without the need for purification. In control experiments, we characterized the binding affinity in buffer and lysate using the conventional method of prescribing different concentrations and determined a low-picomolar *K*_D_ for the antibody. We next performed a time series measurement of the production of GFP in cell-free lysate in the presence of the antibody, thus successfully applying our method to obtain the *K*_D_. Finally, we demonstrate that these measurements can be performed in different sample chamber setups, specifically inside microcapsules, thus potentially substantially increasing the throughput.

## RESULTS

### FCS tracks real-time production of GFP in cell-free lysates

To investigate the ability of FCS to measure fluorescent proteins in a non-pure environment, we measured over several hours the expression of GFP in two *E. coli* derived cell-free lysates, one commercial (Rabbit Biotech) and one homemade (Figure 2). While both lysates alone without plasmid had signal above that of water or buffer, the signals did not show meaningful correlation (Figure 2bc). Interestingly, the photon counts from both lysates decreased in the initial couple of hours before steadily increasing in the commercial lysate while remaining constant in the homemade.

**Figure 2.**
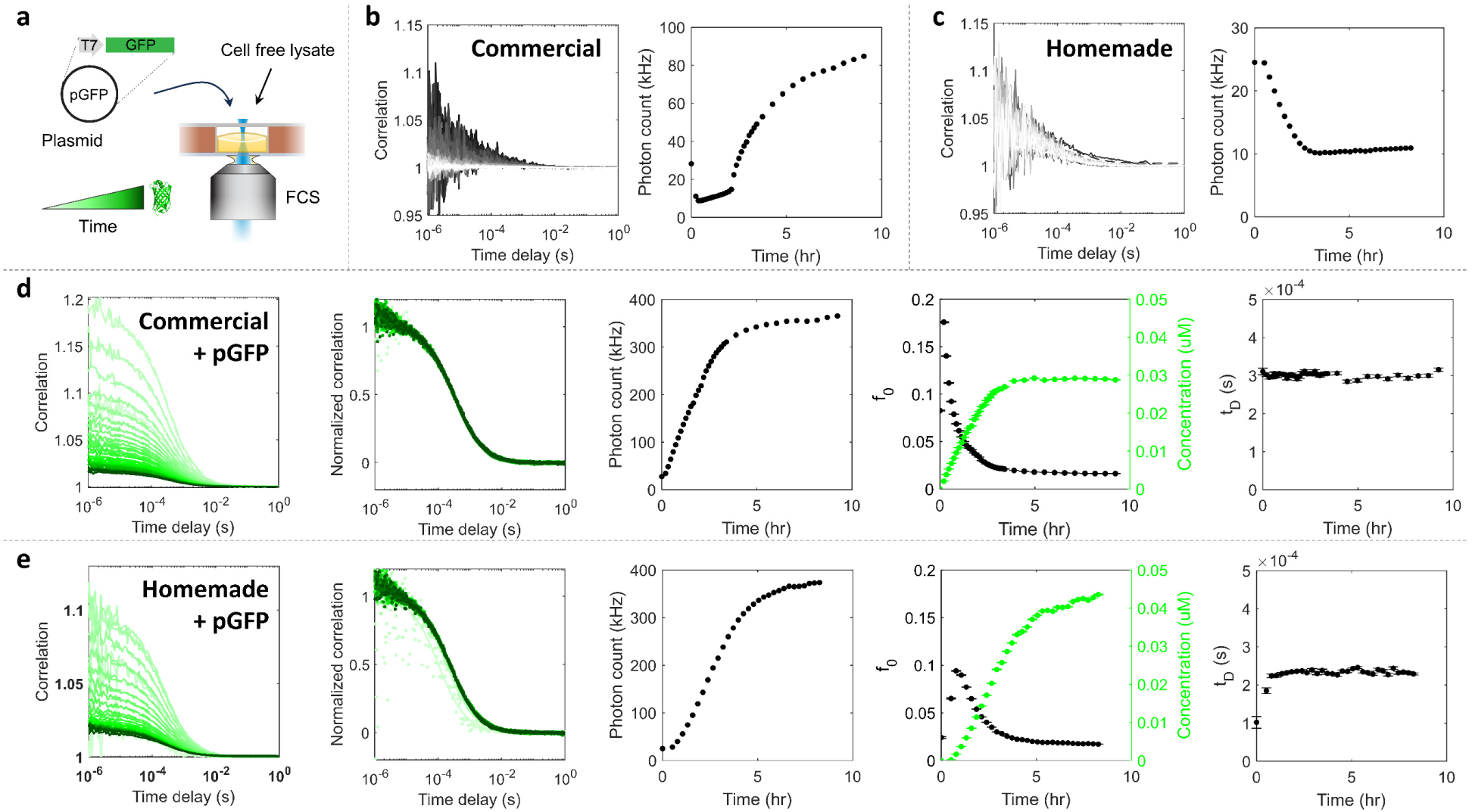
FCS tracks fluorescent proteins synthesized in cell-free lysates in real time. **a.** Production of GFP by cell-free reaction containing a GFP plasmid (pGFP) was measured by FCS over time. **b-c.** Measurements of a commercially available (Rabbit Biotech) **(b)** and homemade **(c)** lysates without plasmid. Left: correlation curves (left) are colored black (earlier time points) to white (later) and show noise from the lysate background. Right: photon counts for both lysates initially decreased before increasing, and the homemade lysate had lower lysate background signal. **d-e.** Measurements of GFP production over time in commercial **(d)** and homemade **(e)** lysates containing 1 µg/ml (0.23 nM) plasmid at room temperature. From left to right: correlation curves, both raw and normalized, colored light (earlier) to dark (later) green; photon count rate; fitted amplitude *f*_0_ and diffusion times *τ*_D_ (Equation 1) over experimental time. Concentrations (green) were calculated from the *f*_0_ given the calibrated detection volume and were corrected for background signal using the lysate-only data at every time point (Equations 2 and 3). The initially lower *f*_0_ and *t*_D_ are due to background noise contribution at low GFP concentrations.

Above the lysate background, fluorescent signal from the GFP protein was detectable in the low nanomolar range within half an hour of plasmid addition (Figure 3de). The *g*^(2)^ correlations as a function of time delay *τ* were fitted to a simple one-species 2D diffusion model

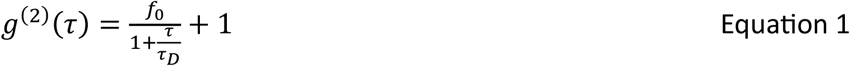

to obtain the amplitude *f*_0_ and diffusion time *τ*_*D*_ corresponding to the average time a molecule takes to traverse the focal volume. The measured concentration *C* at every time point was calculated from the fitted amplitude given the calibrated focal volume *V* and corrected for background contribution using a correction factor χ^2^:

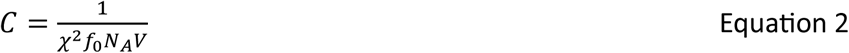

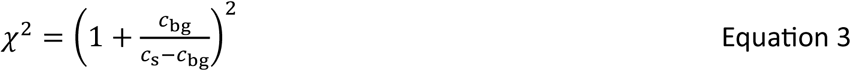

where *N*_*A*_ is Avogadro’s number, *c*_bg_ is the background photon count rate measured by the lysate-only sample at the corresponding time point, and *c*_s_ is the count rate of the sample ^16^. As GFP was produced in the lysates, photon counts increased, *f*_0_ decreased, measured concentration increased, and *τ*_*D*_ remained constant, as expected. The constant diffusion time was also apparent in the normalized plots of correlations. When FCS measurements were started early enough to capture the earliest time points, the total signal from these early points had dominant contribution from the lysate background, resulting in noisier correlations with lower amplitudes (Figure 3e). This explains the initial increase in fitted *f*_0_ as the amount of GFP increased above the background, as well as the initial lower fitted *τ*_*D*_.

**Figure 3.**
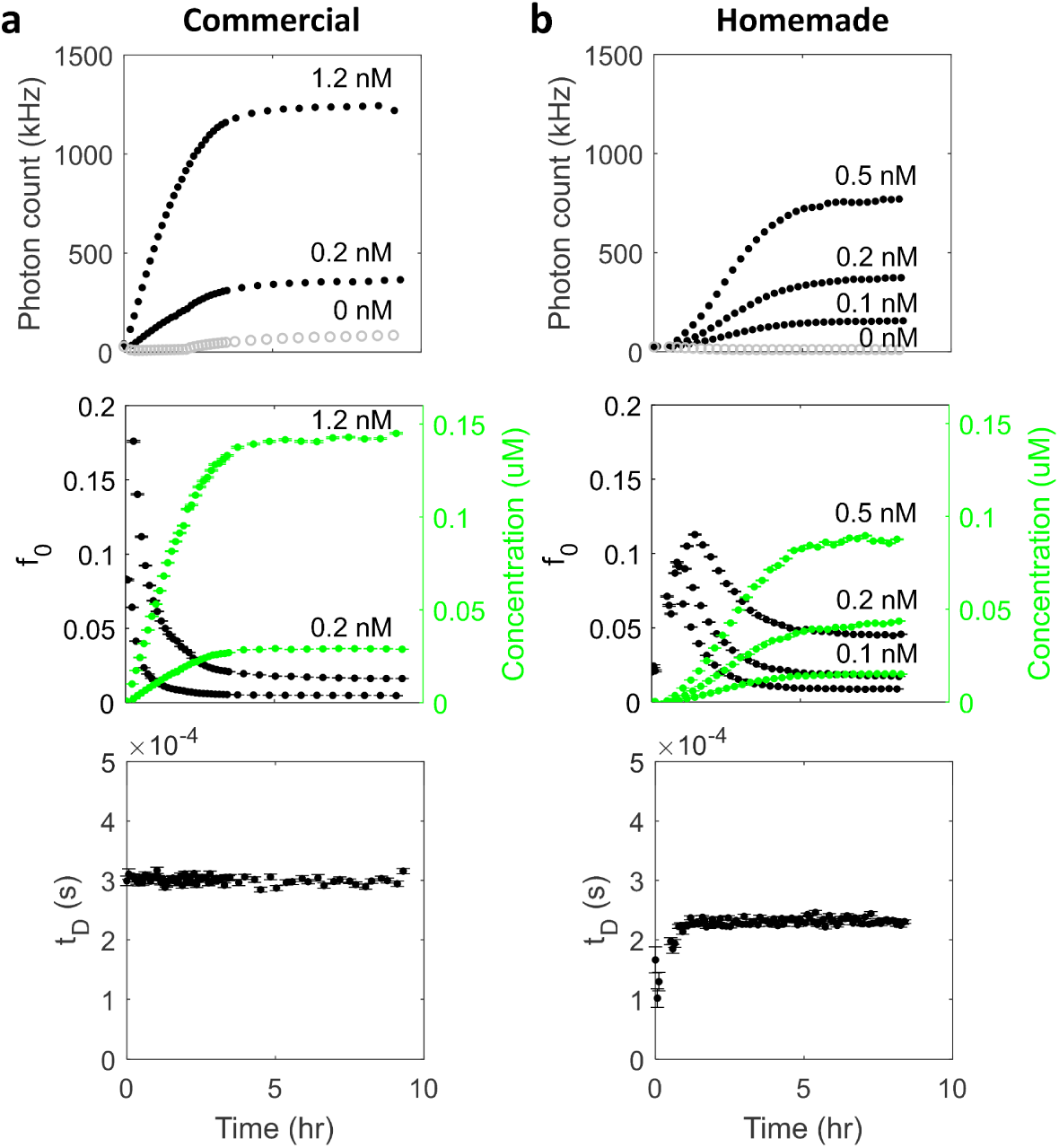
Rate and amount of protein synthesis can be tuned by plasmid concentration. **a-b.** Photon count rate (top), fitted amplitude *f*_0_ (middle) and diffusion times *t*_D_ (bottom) are plotted over experimental time for several GFP plasmid concentrations in commercial (1.2 and 0.2 nM) **(a)** and homemade (0.1, 0.2, and 0.5 nM) **(b)** lysates. Lysate-only photon count rates are plotted as gray open circles. Calculated concentrations included background subtraction. The homemade lysate had slightly slower production rates than the commercial lysate but similar final yields.

Production of GFP was comparable in the commercial and homemade lysates. The experiment shown used 1 µg/ml (0.23 nM) plasmid in both lysates and were performed at room temperature, yielding ∼30-40 nM protein by ∼5 hr. Interestingly, GFP diffusion times were consistently higher in the commercial lysate (*τ*_*D*_ ∼0.3 ms) than in homemade (*τ*_*D*_ ∼0.2 ms). Since *τ*_*D*_ measured in the homemade lysate was the same as that measured in buffer, the higher *τ*_*D*_ in the commercial lysate must be due to certain conditions or components of the lysate.

To summarize, FCS was demonstrated to successfully measure the production of GFP starting from single digit nanomolar concentrations in the unpurified environment of cell-free lysates. In comparison to the commercial lysate, our homemade lysate produced similar or even higher protein while its background remained low.

### Tuning the rate and amount of protein synthesis by plasmid concentration

Accurate measurements of protein binding affinity require appropriate regimes of the concentrations of binding partners ^17^. Ideally, the concentration of one binding partner (the “trace” species) should be much lower than the expected *K*_D_ while the concentration of the other partner should be varied to capture the full range of fraction of the trace species bound. In any assay, a practical constraint to how low in concentration the trace species can be, as needed to measure very low *K*_D_s, is the lowest detectable signal above background. While picomolar concentrations of GFP (and thus *K*_D_s) are measurable in our FCS system when using buffer, the detection limit rises to low nanomolar in the background of cell-free lysate, thus also raising the lowest measurable *K*_D_s to the low nanomolar range. Furthermore, in our approach to using time resolved protein synthesis as the concentration variable, the rate and amount of synthesis must be tuned to achieve the full range of fraction bound. If the protein is made too quickly, the low-concentration region of the binding curve may not be captured, whereas if the protein is made too slowly, the cell-free reaction may stop before reaching a final yield high enough to capture the high concentration region.

Thus, to optimize the synthesis rate and yield for binding measurements and to further characterize the commercial and homemade lysates, we measured the production of GFP using different amounts of plasmid (Figure 3). While much higher plasmid concentrations (∼10-15 µg/ml, or 2.3-3.5 nM) are typically used in cell-free reactions to achieve high yield, we tested a lower range (0.5-5 µg/ml, or 0.12-1.2 nM) because slower protein synthesis better captures the low concentrations required for measuring the expected low *K*_D_ between GFP and its antibody. As expected, higher plasmid concentrations resulted in faster rates and higher final yields. 5 µg/ml plasmid produced ∼140 nM GFP within 5 hr at room temperature (Figure 3a). Using 10x less plasmid (0.05 µg/ml in the homemade lysate), our FCS system was still able to measure a clear signal above lysate background with clean correlations, plateauing at ∼15 nM GFP in 5 hr (Figure 3b). Diffusion times of expressed GFP (*τ*_*D*_ ∼0.3 ms and ∼0.2 ms in commercial and homemade lysates, respectively) were independent of plasmid concentration and remained constant over time, as expected. Overall, the homemade lysate had slightly slower production rates than the commercial lysate but similar final yields.

### Binding affinity between GFP and antibody in buffer and lysates

As control experiments, we first characterized the binding affinity between GFP and its antibody in buffer and lysate using the conventional method of prescribing different concentrations. When the concentration of GFP was held constant (200 pM) and the concentration of antibody was titrated in buffer (Figure 4a), correlation curves shifted rightward to higher overall diffusion times when fitted to the one-species model (Equation 1) (Figure 4bc). Correlations were then fitted to a 2-species model:

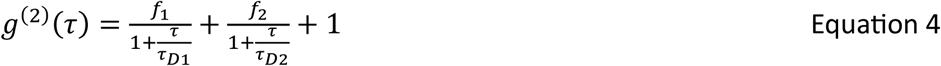

**Figure 4.**
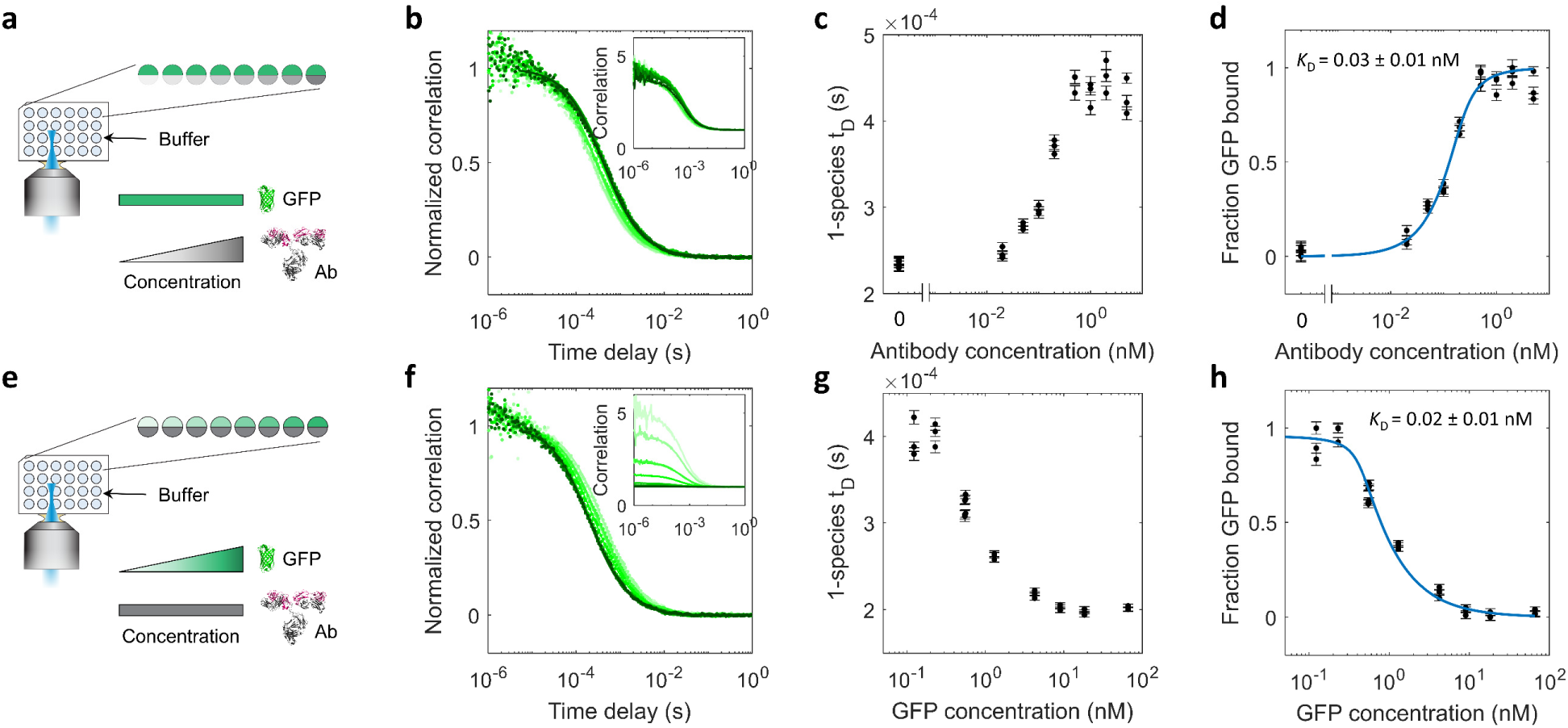
Binding affinity between GFP and an anti-GFP antibody in buffer solution. **a.** To measure the binding affinity between GFP and an anti-GFP antibody, samples containing a constant concentration of GFP (200 pM) and varying concentrations of the unlabeled antibody in buffer were measured by FCS. **b.** Normalized correlation curves shifted right at higher antibody concentrations. Inset shows non-normalized correlations. **c.** Diffusion times from one-species fits (Equation 1) to correlations increased as a function of antibody concentration. **d.** Fraction of GFP bound to antibody was calculated from two-species fits to correlations (Equations 4 and 5). The quadratic binding equation (Equation 6) was fitted to the binding data to obtain the dissociation constant *K*_D_ = 0.03 ± 0.01 nM and GFP concentration [*G*] = 0.19 ± 0.03 nM, which agrees with the known amount of GFP used. **e.** A binding curve was also obtained by varying the GFP concentration while keeping constant the unlabeled antibody concentration (400 pM). **f.** Normalized correlation curves shifted left at higher GFP concentrations. Inset shows non-normalized curves with lower amplitudes at higher GFP concentrations. **g.** Diffusion times from one-species fits decreased as a function of GFP concentration. **h.** Fraction of GFP bound calculated from two-species fits to correlations, plotted against GFP concentration. The variant quadratic binding equation (Equation 7) was fitted to the measurements to obtain *K*_D_ = 0.02 ± 0.01 nM and antibody concentration [*a*] *=* 0.41 ± 0.02 nM, consistent with both the *K*_D_ determined by the standard binding method **(a-d)** and the known amount of antibody used. This antibody and GFP have very tight binding with low picomolar *K*_D_.

where *f*_1_ and *f*_2_ correspond to the amplitudes in each population with diffusion times *τ*_*D*1_ and *τ*_*D*2_. *τ*_*D*1_ represents the diffusion time of GFP alone, and *τ*_*D*2_ represents that of GFP bound to antibody. The fraction of GFP bound was then calculated by

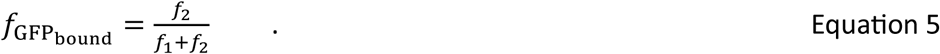

The calculated fraction GFP bound increased through the range of ∼10 – 500 pM antibody and saturated above ∼500 pM (Figure 4d), suggesting that the *K*_D_ was in the picomolar range. Since the concentration of the “trace” or limiting component (200 pM GFP) was not << *K*_D_, the data was not fitted to the simple hyperbolic binding equation, and *K*_D_ could not be simply read as the concentration of antibody at 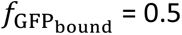, but rather was determined from a fit to the quadratic binding equation:

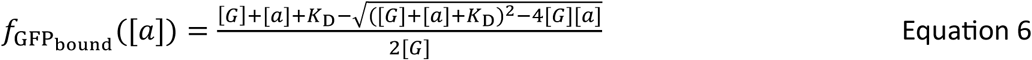

where [*G*] is the total concentration of the trace component which was held constant (GFP), [*a*] is the total concentration of the titrated component (antibody), *K*_D_ is the dissociation constant, and the equation gives the fraction of GFP that is bound to antibody. Equation 6 was fitted to the measured binding curve (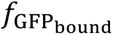 vs. antibody concentration [*a*]) with two fitting parameters [*G*] and *K*_D_. From the fit, GFP and its antibody had *K*_D_ = 0.03 ± 0.01 nM, and [*G*] = 0.19 ± 0.03 nM, which agrees well with the known 200 pM GFP used.

A binding curve could also be obtained by titrating the concentration of GFP while keeping constant that of the unlabeled antibody (Figure 4e). This case mimics the proposed cell-free expression approach in which the fluorescently labeled component increases in concentration over time. Since FCS measures the fluorescent species, as GFP is produced and rises in concentration in excess of the 400 pM antibody used, correlation curves shifted leftward to lower overall diffusion times (Figure 4fg). Correlations were fitted to the 2-species model (Equation 4), from which the fraction of GFP bound was calculated (Equation 5) and plotted against GFP concentration (Figure 4h). This binding curve was then fitted to a variation of the quadratic binding equation which gives the fraction of GFP bound to antibody as a function of varying the concentration of GFP:

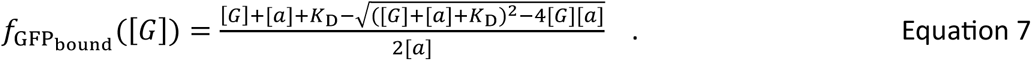

Equation 7 was fitted to the measured binding curve (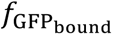 vs. GFP concentration [*G*]) and yielded *K*_D_ = 0.02 ± 0.01 nM and antibody concentration [*a*] = 0.41 ± 0.02 nM, consistent with both the *K*_D_ determined by the standard titration method (Figure 4a-d) and the known amount of antibody used.

We next repeated the control binding measurements in cell-free lysates (Figure 5). Higher amounts of the non-varying protein were required than in the buffer experiments in order to measure GFP signals above the higher background due to lysate. In commercial lysate, when GFP was held constant at 2 nM and the concentration of antibody was titrated, fitting the fraction of GFP bound to Equation 6 produced *K*_D_ = 0.04 ± 0.02 nM and [*G*] = 1.9 ± 0.1 nM (Figure 5b), consistent with the *K*_D_ measured in buffer and the amount of GFP used. In homemade lysate, measurements using 5 nM GFP and varying the concentration of antibody produced *K*_D_ = 0.25 ± 0.17 nM and [*G*] = 5.3 ± 0.6 nM (Figure 5c). While [*G*] was accurately determined, the measured apparent *K*_D_ was higher than that measured in buffer and had greater uncertainty due in part to the use of GFP at a concentration much greater than *K*_D_.

**Figure 5.**
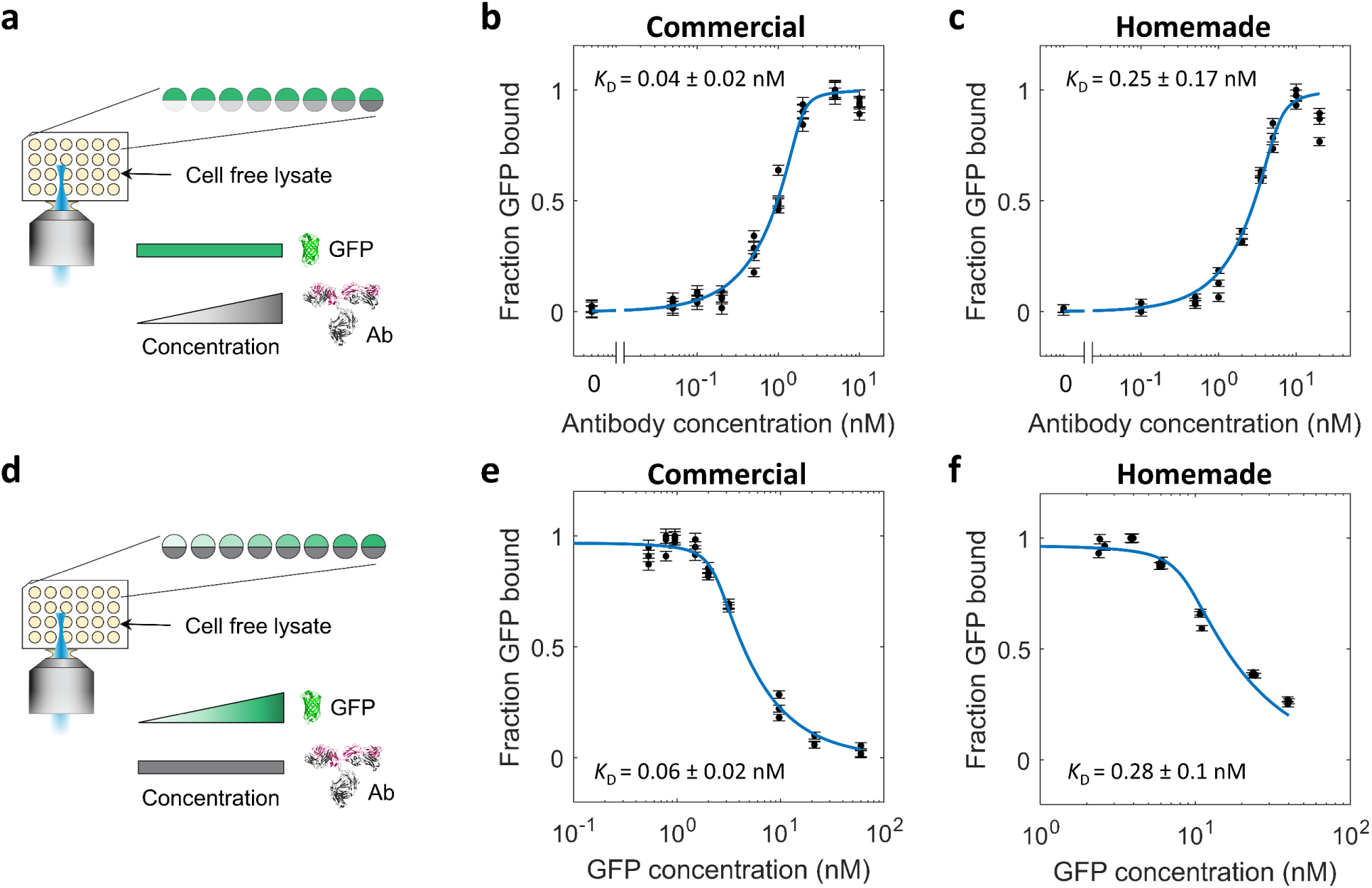
Binding affinity between GFP and an anti-GFP antibody in cell-free lysates. Control experiments as described in Figure 4, varying either unlabeled antibody **(a-c)** or GFP **(d-e)**, were repeated in cell-free lysates. Higher amounts of the non-varying protein were required than in the buffer experiments in order to measure GFP signals above the higher background due to lysate. **(b)** Vary antibody in commercial lysate, using 2 nM GFP. Fitted *K*_D_ = 0.04 ± 0.02 nM, GFP concentration [*G*] = 1.9 ± 0.1 nM. **(c)** Vary antibody in homemade lysate, using 5 nM GFP. Fitted *K*_D_ = 0.25 ± 0.17 nM, GFP concentration [*G*] = 5.3 ± 0.6 nM. **(e)** Vary GFP in commercial lysate, using 2 nM antibody. Fitted *K*_D_ = 0.06 ± 0.02 nM, antibody concentration [*a*] = 2.2 ± 0.1 nM. **(f)** Vary GFP in homemade lysate, using 10 nM antibody. Fitted *K*_D_ = 0.28 ± 0.1 nM, antibody concentration [*a*] = 8.1 ± 0.4 nM.

When GFP concentration was varied while antibody concentration was kept constant, the fraction of GFP bound was measured as a function of GFP concentration and fitted to Equation 7. In commercial lysate using 2 nM antibody, we obtained *K*_D_ = 0.06 ± 0.02 nM and [*a*] = 2.2 ± 0.1 nM (Figure 5e). In homemade lysate using 10 nM antibody, we obtained *K*_D_ = 0.28 ± 0.1 nM and [*a*] = 8.1 ± 0.4 nM (Figure 5f). Again, while [*a*] was accurately determined in both lysates, the measured apparent *K*_D_ was higher in the homemade lysate due in part to the higher antibody concentration used which precludes accurate determination of the very low *K*_D_. In addition, it is also possible that the binding affinity between the GFP and the antibody was slightly altered in the conditions of the homemade lysate buffer.

In summary, we measured the binding affinity of GFP and its antibody by titrating either GFP or antibody and found good agreement between the two methods that both yielded a picomolar *K*_D_.

### Determination of binding affinity from FCS measurements of protein synthesis over time in cell-free lysates

Samples containing GFP plasmid and antibody in both cell-free lysates were measured over time by FCS for production of GFP and binding to antibody (Figure 6a). In a ∼6 µl solution containing 5 µg/ml (1.2 nM) plasmid and 20 nM antibody in commercial lysate, GFP was quickly produced and reached ∼200 nM within 2 hours (Figure 6b,c). Further data beyond 2 hr was not collected because the amount of GFP reached 10x higher than the antibody concentration and was sufficient for a binding curve. Normalized correlation curves shifted left as free GFP was produced whose concentration increased above that of the antibody (Figure 6b). The total GFP concentration at each time point was determined from the total amplitude *f*_0_ = *f*_1_ + *f*_2_ of two-species fits (Equation 4) and background subtracted (Equations 2 and 3) using lysate-only data (Figure 6c). The binding curve results from plotting the fraction of GFP bound calculated from two-species fits (Equations 4 and 5) against the measured GFP concentrations (Figure 6d). The variant quadratic binding equation (Equation 7) was then fitted to the measurements to obtain *K*_D_ = 11 ± 1.5 nM and [*a*] = 19 ± 1.3 nM. This process was repeated in homemade lysate using 2 µg/ml (0.5 nM) plasmid and 20 nM antibody and yielded *K*_D_ = 15 ± 3 nM and [*a*] = 20 ± 2 nM (Figure 6e-g). In both lysates, the fitted antibody concentration [*a*] matched the known concentration used (20 nM), while the *K*_D_ ∼10 nM was much higher than the picomolar affinity determined in control experiments.

**Figure 6.**
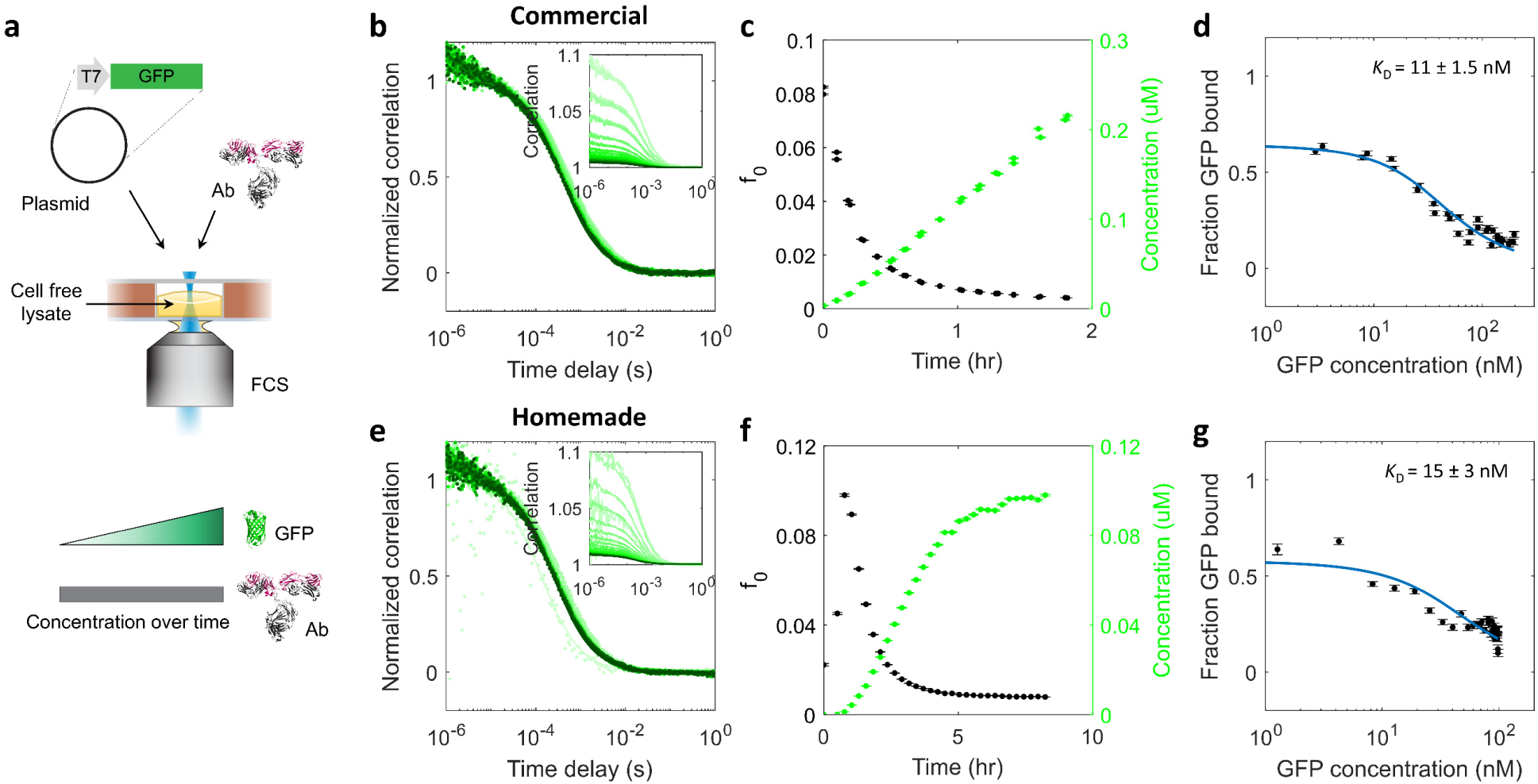
Determination of binding affinity from FCS measurements of protein synthesis over time in cell-free lysates. **a.** Samples containing GFP plasmid and antibody in cell-free lysate were measured over time by FCS for production of GFP and antibody binding. **b-d.** 5 µg/ml (1.2 nM) plasmid and 20 nM antibody in commercial lysate. Normalized correlation curves **(b)** shifted left as GFP concentration increased over time (**b** inset). Total GFP concentrations were determined from the total amplitude *f*_0_ of two-species fits (Equation 4) and background subtracted (Equations 2 and 3) using lysate-only data at each time point **(c)**. Fraction of GFP bound calculated from two-species fits (Equations 4 and 5) is plotted against the measured GFP concentrations **(d)**. The variant quadratic binding equation (Equation 7) was fitted to the measurements to obtain *K*_D_ = 11 ± 1.5 nM and antibody concentration [*a*] = 19 ± 1.3 nM. **e-g.** Correlation curves **(e)**, total amplitudes and background-subtracted concentrations **(f)**, and binding curve **(g)** for measurements of 2 µg/ml (0.5 nM) plasmid and 20 nM antibody in homemade lysate. Note that the amplitude was initially low **(f)** due to the dominant contribution of lysate background to the total signal, as also evident in the earliest correlations **(e)**. Fitted *K*_D_ = 15 ± 3 nM, antibody concentration [*a*] = 20 ± 2 nM. The fitted antibody concentrations are consistent with the known amount (20 nM) used. Since the concentration of antibody, required to be sufficiently high to measure signal above lysate background, was much higher than the true low-picomolar *K*_D_ (Figures 4 and 5), these real-time binding curves establish an upper bound on the *K*_D_.

The higher measured *K*_D_s resulted from the relatively high antibody concentration used and from reasons discussed in the next section. The high antibody concentration was necessary for measurements above lysate background (as discussed previously in the section “Tuning the rate and amount of protein synthesis by plasmid concentration”) but precluded accurate determination of the low picomolar *K*_D_. These higher measured *K*_D_s therefore establish an upper bound on the true binding affinity. The ability to measure a low-nanomolar *K*_D_ is still very informative because weak and strong binders would be distinguishable in a large initial screen and because most biological affinities range from nanomolar to micromolar. Importantly, we have demonstrated that this determination of *K*_D_ is possible within a couple hours of starting a one-pot cell-free reaction.

### Experimental factors affecting determination of binding affinity

It is known that accurate determination of binding affinity is not possible in the presence of experimental noise when the concentration of the trace species is high compared to the *K*_D_, leading to a measured value greater than the true *K*_D_ ^17^ (Supplementary Figure 2). To test that this explains the much higher apparent *K*_D_s (∼10 nM) measured by tracking real-time protein synthesis (Figure 6) in comparison to the true *K*_D_ (∼30 pM) (Figures 4 and 5), and to characterize the variability in the method, we performed multiple cell-free production experiments using different concentrations of the “trace species” (antibody [*a*]). A range of antibody concentrations from [*a*] ∼3 nM to 80 nM was used. Note that the lowest [*a*] used was still much higher than the picomolar *K*_D_ because of the constraint due to lysate background: a much lower picomolar [*a*] would have meant that by the time the GFP signal was measurable above the lysate background, at a few nanomolar GFP, almost all of the GFP would have been free, as the case for the remainder of the time series, thus yielding an uninformative null binding curve.

We found that the measured *K*_D_ reproducibly ranged between ∼2-20 nM across all experiments (open black markers in Figure 7a). While these values were ∼100-500x higher than the picomolar *K*_D_s measured in control experiments performed without real-time protein production (solid black markers in Figure 7a), they provided a confident low-nanomolar upper bound. However, surprisingly, the measured *K*_D_s were independent of the antibody concentration. At [*a*] ∼10 nM and below, *K*_D_s measured by protein synthesis were still ∼10-50x higher than those measured by control experiments at similar [*a*]’s. Thus, the use of [*a*] >> *K*_D_ could not be the only factor causing the higher measured *K*_D_s.

**Figure 7.**
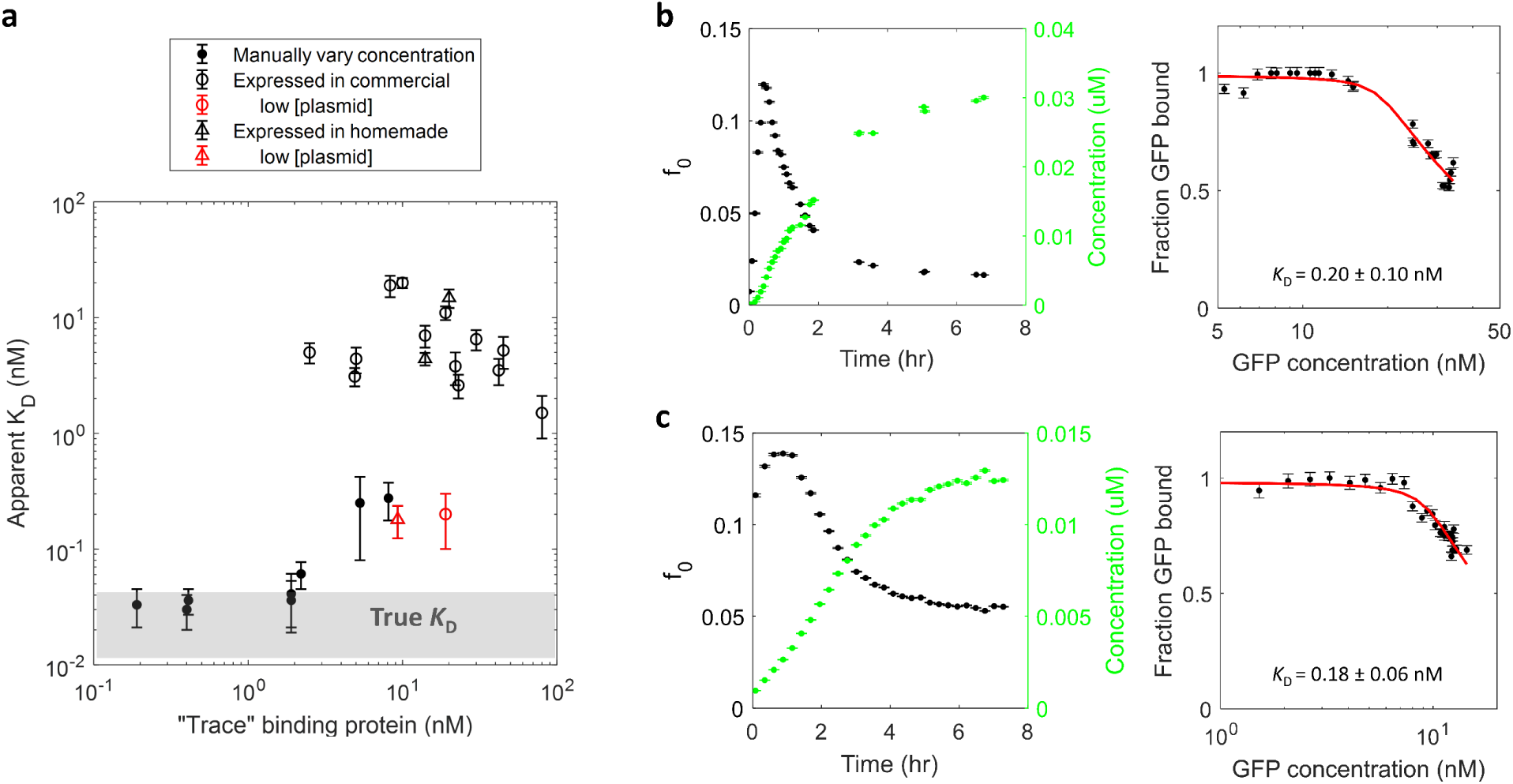
Use of higher concentrations of trace binding partner and other experimental factors establish upper bound on the *K*_D_ measured by cell-free protein synthesis. a. Compilation of the measured *K*_D_ vs. the “trace” protein concentration *P* used from all binding experiments. Accurate determination of *K*_D_ typically requires *P* < *K*_D_, a condition often not met in biological experiments due to practical constraints. Control experiments, in which sub-nanomolar *P* (antibody or GFP concentration) were used (solid markers) (from Figures 4 and 5), suggested that the true affinity was ∼30 pM or less (shaded area). Much higher *K*_D_s were measured from real-time protein synthesis experiments (open markers) due to a combination of the much higher *P* (antibody concentration) required in lysate background, GFP maturation time, and/or nonequilibrium binding. Slowing down and lowering protein expression produced *K*_D_ measurements more consistent with control experiments (red open markers) **(b, c)**. **b.** Binding experiment from protein synthesis using 0.5 µg/ml (0.12 nM) GFP plasmid and 20 nM antibody in commercial lysate. Left: total GFP concentrations were determined from the total amplitude *f*_0_ of two-species fits (Equation 4) and background subtracted (Equations 2 and 3) using lysate-only data at each time point. Right: fraction of GFP bound calculated from two-species fits (Equations 4 and 5) is plotted against the measured GFP concentrations. The variant quadratic binding equation (Equation 7) was fitted to the measurements to obtain *K*_D_ = 0.2 ± 0.1 nM and antibody concentration [*a*] = 18.7 ± 0.5 nM (red open circle in **(a)**). **c.** Total amplitudes and background-subtracted concentrations (left), and binding curve (right) for measurements of 0.5 µg/ml (0.12 nM) plasmid and 20 nM antibody in homemade lysate. Fitted *K*_D_ = 0.18 ± 0.06 nM, antibody concentration [*a*] = 9.3 ± 0.2 nM (red open triangle in **(a)**).

Two additional factors may be the maturation time of GFP and nonequilibrium binding. First, maturation times of fluorescent proteins have been found to range from several minutes to upwards of an hour ^18^, before which the protein is folded but non-fluorescent. During this time in a protein synthesis experiment, dark GFP molecules would have been able to bind the antibody but would not have been detected by FCS, thus sequestering a fraction of the antibody to be unavailable to bind to the matured bright GFP molecules. This would result in a lower apparent fraction of GFP bound, as measured by bright GFP molecules, and thus a higher apparent *K*_D_. Second, proteins that have very low *K*_D_s may have very slow binding on rates ^17^. In our experiments that measured binding during 2-5 hrs of GFP synthesis, the production and detection of GFP might have outpaced the on rate or time required for equilibrium binding, resulting again in a lower apparent fraction of GFP bound and higher apparent *K*_D_.

To test both factors, we slowed down the rate of protein synthesis by using a lower GFP plasmid concentration (0.5 µg/ml) in binding experiments. GFP production rate was ∼7 nM/hr in the commercial lysate (Figure 7b) and ∼2.5 nM/hr in the homemade lysate (Figure 7c), compared to ∼100 nM/hr (Figure 6c) and 20 nM/hr (Figure 6f) using 5 µg/ml and 2 µg/ml previously in the two lysates, respectively. We found that indeed the measured *K*_D_ was lowered: *K*_D_ = 0.2 ± 0.1 nM in the commercial lysate, *K*_D_ = 0.18 ± 0.06 nM in the homemade lysate (Figure 7bc). By slowing down protein expression, these *K*_D_ measurements were now consistent with control experiments done at similar antibody concentrations (red open markers vs. black solid markers in Figure 7a). Furthermore, in the presence of experimental noise, these binding curves corresponding to ∼200 pM *K*_D_s are indistinguishable from those corresponding to ∼30 pM when using excess (20 nM) antibody (Supplementary Figure 2), which suggests that now the *K*_D_ determination is limited by the high trace protein concentration.

### Determination of binding affinity from protein synthesis inside microcapsules

Towards high throughput screening applications, our FCS-based in-lysate binding method may be adapted to perform measurements in various sample formats including well plates, microfluidic chambers, and microcapsules. The method is readily adaptable for measurements in plates and microfluidic chips that have a glass bottom compatible with the imaging objective. Microencapsulation provides a method to compartmentalize thousands to millions of independent reactions. Through a double emulsion droplet generation process using a homemade microfluidic device, we packaged cell-free reactions into 200-300 µm droplets enclosed by a polymer shell, which are then stabilized by UV crosslinking of the polymer (Figure 8a). Each capsule contains only ∼10 nL of solution. It has not been shown in the literature whether cell-free reaction can proceed inside microcapsules or whether the fluorescence correlation signal can be measured through the capsule shell.

**Figure 8.**
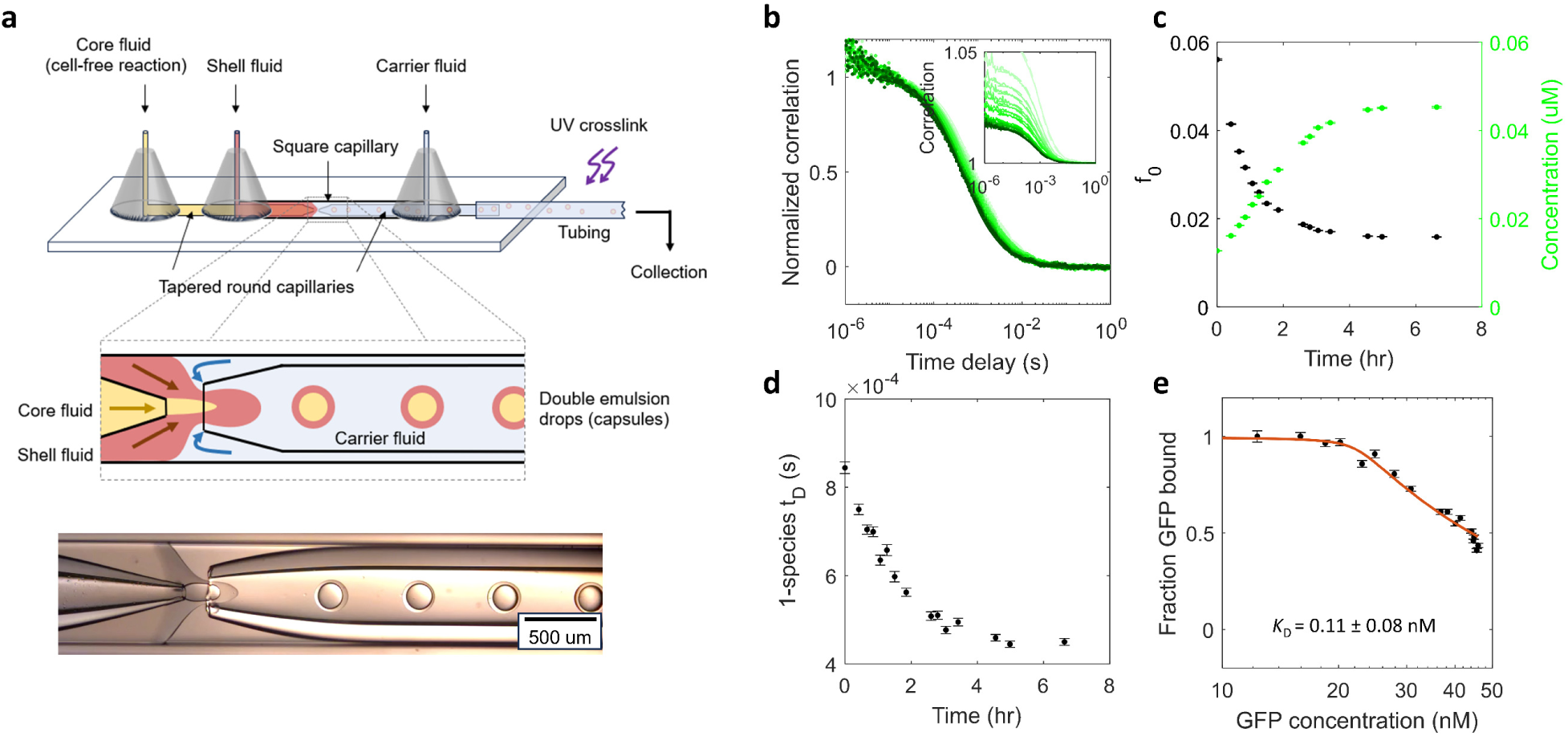
Determination of binding affinity from protein synthesis inside microcapsules. **a.** Top: schematic of double emulsion droplet generation on a microfluidic device. Through the flow of fluids in two round capillaries with tapered openings within a larger square capillary, the core fluid (cell-free reaction) (yellow) is packaged into droplets with a polymer shell (red) surrounded by the carrier fluid (light blue). Each capsule contains ∼10 nl or less of reaction. Droplets pass under a UV lamp to crosslink the shell polymer before collection and measurement of the capsules by FCS. Bottom: image of droplet formation in the region of the interface between the three fluids. **b-e.** FCS measurements of protein synthesis and binding inside a microcapsule containing 1.45 µg/ml (0.34 nM) GFP plasmid and 20 nM antibody in commercial lysate. **b.** Normalized correlation curves shifted left as GFP concentration increased over time (inset). **c.** Total GFP concentrations were determined from the total amplitude *f*_0_ of two-species fits (Equation 4) and background subtracted (Equation 2 and 3) using lysate-only data at each time point. **d.** Diffusion times from one-species fits decreased over time as GFP concentration increased. **e.** Fraction of GFP bound calculated from two-species fits (Equations 4 and 5) is plotted against the measured GFP concentrations. The variant quadratic binding equation (Equation 7) was fitted to the measurements to obtain *K*_D_ = 0.11 ± 0.08 nM and antibody concentration [*a*] = 22.2 ± 0.5 nM.

To investigate the ability to perform FCS measurements through the polymer shell, we made microcapsules containing cell-free lysate, GFP plasmid, and GFP antibody, placed the microcapsules on a glass coverslip, and measured the FCS signal inside for several hours. We were able to measure both protein synthesis and binding consistent with previous measurements made in wells. Inside a capsule containing 1.45 µg/ml (0.34 nM) GFP plasmid and 20 nM antibody in commercial lysate, GFP concentration increased to ∼45 nM in ∼4 hours, while diffusion times from one-species fits decreased (Figure 8b-d). The fraction of GFP bound was calculated from two-species fits (Equations 4 and 5) and plotted against the measured GFP concentrations to yield the binding curve (Figure 8e). A fit of the variant quadratic binding equation (Equation 7) gave *K*_D_ = 0.11 ± 0.08 nM and antibody concentration [*a*] = 22.2 ± 0.5 nM, both agreeing with the previous binding measurements and known amount of antibody used. In summary, real-time FCS measurements of proteins synthesized by cell-free lysate enable binding affinity determination, and this method can be used on a variety of sample formats including microencapsulation for high throughput screening applications.

## DISCUSSION

Relying on cell-free protein synthesis has both limitations and advantages. A fundamental requirement is that the protein must be amenable to expression, and to a high enough extent, in one of the ∼10-20 existing cell-free platforms ^13^. But this may also be a challenge in cell-based systems for difficult-to-synthesize proteins. In fact, some proteins that do not express well in cells due to toxicity or aggregation may be better tolerated and produced in cell-free systems.

For example, soluble membrane proteins may be made co-translationally in lipid nanodiscs by cell-free systems ^19,20^. Furthermore, for proteins that do not purify well, the ability to measure binding in the lysate environment bypasses the purification requirement. Finally, tracking protein synthesis by FCS provides a wealth of information in addition to binding affinity: expression levels, rates, and aggregation tendencies, all valuable information for screening new protein variants. All of this information is obtained with minimal amounts of cell-free reaction: ∼6 µl for reactions in wells, and less than 10 nl inside a microcapsule.

Although measuring binding simultaneous with protein synthesis eliminates the need for protein purification and significantly speeds up affinity assessment, this strategy has inherent limitations. First, without purification, the lowest detectable limit of the expressed protein is ∼1 nM to be above the lysate background. Second, binding may not reach equilibrium since measurements are made during continuing protein synthesis. Third, the fluorescent protein used to label the protein of interest has a non-negligible maturation time, which obscures readout of the true protein concentration. In this work, the choice of using GFP and its antibody as the model pair, with their low-picomolar *K*_D_, enabled us to characterize these limitations. We found that these three factors limit the lowest measurable *K*_D_ to the low nanomolar range, a sensitivity sufficient for many screening applications, nonetheless.

We found that these limitations may be mitigated by slowing the rate of protein synthesis, allowing the measurement of sub-nanomolar *K*_D_ (Figure 7). The rate can be controlled by adjusting the plasmid concentration and/or temperature. To mitigate complications arising from GFP maturation time, we kept a layer of air above the thin layer of cell-free solution. Future improvements may include saturating the reaction with oxygen or supplying higher oxygen-content air above the solution. An alternative strategy may employ a split-GFP technique in which a pre-matured GFP1-10 is added to the cell-free reaction to complement the protein-GFP11 being expressed, since pre-maturation has been shown to significantly speed up signal detection ^21^. Finally, to avoid the issue of fluorescent protein maturation entirely, chemical dyes may be used as the fluorescent label instead of fluorescent proteins. Specifically, BODIPY-FL-labeled lysine (Promega Corporation) may be incorporated into the protein during translation, and the free lysine-dye component in the correlation data can be removed during analysis.

While a shift in diffusion time of the FCS correlation curves provides a simple readout of binding requiring a simple one-color FCS setup, the diffusion times of the two species (unbound and bound) should differ by at least 1.6x to be distinguishable under typical conditions ^22^. This requirement means that the masses of the two species must differ by at least 4x, since diffusion time is inversely proportional to the radius (cubed root of mass). In the current work, FCS easily and reliably measured the increase in diffusion time of GFP upon antibody binding based on ∼6x increase in mass, from ∼30 kDa GFP to ∼30+150 kDa GFP+antibody. Measurements of other binding pairs that have smaller changes in size can be made possible using dual-focus FCS, which can measure Angstrom-scale changes ^23,24^, or dual-color FCS, in which binding is indicated by cross-correlation between proteins labeled with different dyes ^25^ ^7^.

By combining cell-free protein synthesis and FCS, we simultaneously perform the synthesis and affinity measurement steps, thus enabling a simpler, faster workflow for screening. Suppose we need to measure whether variants of a protein bind to another protein of interest, for example in a screen for antibody fragments targeting the SARS-CoV-2 spike protein. Conventionally, each variant must be separately expressed in cells and purified. Then, for each variant, multiple samples containing different concentrations of the protein must be prepared and measured to obtain a binding curve. This process typically takes several days, from protein expression to purification to measurements. In contrast, our approach measures production and binding in a one-pot reaction for each variant without the need for purification or prescription of different concentrations, using only a few hours of cell-free protein synthesis (Supplementary Figure 3). This significantly simplifies and speeds up measurements, enabling scalability and high throughput in screens for biomolecular binding.

We have demonstrated the proof of concept on performing simultaneous measurements of protein production and binding, leaving potential for extensive future technology development for high throughput applications. In a high throughput screen, repeated measurements are to be made over time on multiple samples to track protein production and binding (Supplementary Figure 3). Currently each measurement takes 30 sec on the conventional FCS, which enables thousands of measurements in a day. By using multi-spot FCS ^26^ or selective plane illumination FCS (SPIM-FCS) ^27^ coupled to single photon avalanche diode (SPAD) array detectors ^28^, tens to thousands of measurements can be made in parallel, thus reducing measurement times to < 1 s and enabling potentially many thousands of samples to be measured in one day. This ability presents a truly revolutionary potential in high throughput studies of intermolecular interactions across biosciences and medicine.

## METHODS

### Preparation of CFPS lysates

Overnight starter cultures of ClearColi BL21(DE3) (Biosearch Technologies) grown in 2x YT media supplemented to 1% NaCl at 37 C 225 rpm were diluted 1:50 into multiple 2 L bafled shake flasks containing 500 mL of 2x YT/1% NaCl media. Cultures were grown until OD600 reached 0.5-0.7 and induced with 1 mM IPTG for 4h. Cells were harvested by centrifugation at 8000 rpm in an SLA-3000 rotor, washed with Buffer A (10 mM Tris base, 14 mM magnesium glutamate, 60 mM potassium glutamate, 1 mM DTT), centrifuged again at 6000 rpm, and resuspended with 1 mL of Buffer A per gram of wet cell mass and frozen at −80 C. Roughly 5 g of wet mass per liter of media is generally obtained.

After thawing, cells were lysed through sonication using a qSonica Q500 probe-tip sonicator with 1/8”-1/4” probes, dependent on sample volume, in 30 s on : 30 s off cycles and 25% amplitude until 3,000 kJ of input energy was reached. Cell suspension should turn a darker shade of brown as lysis increases. An additional 1 mM DTT was added after sonication. Sonicated lysates were transferred into 1.5 mL microcentrifuge tubes and centrifuged at 18,000 rcf for 10 min at 4 C and the supernatant transferred to fresh 1.5 mL microcentrifuge tubes. Lysates were incubated at 37 C for 30 min to perform the run-off reaction and centrifuged again at 10,000 rcf for 10 min at 4 C. Supernatant was transferred to fresh 1.5 mL microcentrifuge tubes before flash freezing and storage at −80 C.

### CFPS reactions

Cell-free protein synthesis (CFPS) reactions (1 mL scale) were prepared in 1.5 mL microcentrifuge tubes by combining lysate (25% final reaction volume), plasmid, and water with the following components: 1.2 mM ATP; 0.86 mM GTP, UTP, and CTP; 34 µg/ml folinic acid; 170 µg/ml E. coli tRNA; 2 mM of each amino acid (-glutamic acid), 0.33 mM NAD; 0.27 mM Acetyl-CoA; 1.5 mM spermidine; 1 mM putrescine; 175 mM potassium glutamate; 10 mM ammonium glutamate; 2.7 mM potassium oxalate; 10 mM magnesium glutamate; and 33 mM PEP. A specified amount of pET-15b plasmid encoding His-Avi-tagged GFP DNA was used for each reaction. Anti-GFP antibody (Abcam 1218) was included at the specified concentrations. CFPS reactions were loaded onto coverslips for FCS tracking or, for microencapsulation, into glass syringes and kept on ice before connecting to syringe pumps and microfluidic device.

### Sample slide preparation

Sample slides consisted of an imaging spacer with wells containing the sample solutions sandwiched by two glass coverslips. In initial tests, an imaging spacer 120 µm in thickness with circular well cutouts (Grace BioLabs #470352) was used. The spacer, which had adhesive on both sides, was first adhered to a glass coverslip. After pipetting 2-3 µl of the sample solution (cell-free reaction) into the center of each 6.35 mm-diameter well, the wells were sealed by adhering a second coverslip over the spacer. The solution formed a ∼3 mm area bounded on the top and bottom by the two coverslips and did not fill up the entire area of the well (Supplementary Figure 1a). The sample slide was placed on the FCS objective for data acquisition. We observed the fluorescent signal in each well had a radial spatial gradient, with high signal near the edge of the solution and low signal near the center (Supplementary Figure 1b). This spatial gradient was only observed for cell-free production of GFP; the signal was uniform for purified GFP protein and fluorescent dyes.

We hypothesized that the radial spatial gradient was due to availability of oxygen because the edge of the cell-free reaction solution was in contact with air left in the wells, and oxygen is required for both cell-free protein production and GFP maturation. Therefore, we switched to a thicker 1.8 mm spacer with 3 mm-diameter wells (Grace BioLabs #666208). To provide uniform accessibility to air, we filled the bottom half of the well with 6 µl cell-free reaction, leaving a layer of air above the solution (Supplementary Figure 1c). Now we found uniform signal in xy and z. We used these thicker spacers for all experiments except when measuring microcapsules, for which we used the thin 120 µm spacers to immobilize the capsules between two coverslips.

Samples were kept on ice before slide preparation, and slides were kept on ice before starting data acquisition on the FCS instrument.

### Microencapsulation device fabrication

The microcapillary device was fabricated based on prior work ^29,30^. Briefly, the base of the device is formed by bridging two 2 by 3-inch glass slides by epoxy and two small glass strips. A round glass capillary (15.24 cm long with an outer diameter of 1.0 mm and inner diameter of 0.580 mm, World Precision Instruments, Sarasota, FL) and a square capillary (with an internal width of 1.0 mm, VitroCom, Mountain Lakes, NJ) compose the main components of the device. The square capillary is glued to the base after being cut to the desired length. The round capillary is centered in a pipette puller (Model P-97, Sutter Instruments, Novato, CA) to decrease its diameter in the center under tension and heat, breaking into two equally tapered capillaries. The tapered glass capillaries were then cleaved to the desired final diameters using a microforge station (Micro Forge MF 830, Narishige, Japan). Typical inner diameters of the capillaries ranged from 20 μm to 300 μm. After cleaning in an ethanol solution with 10 minutes of sonication, tips were treated separately with different silane solutions to change the glass hydrophilicity and hydrophobicity. The inner fluid capillary tip is treated to be hydrophobic so that the aqueous inner fluid could be easily repelled to break up into drops at the end of the capillary. Similarly, hydrophilic coating was applied to the exit capillary to accelerate the breakup of the oil-based middle fluid.

### Microencapsulation

Syringe pumps (Harvard PHD Ultra, Harvard Apparatus, Holliston, MA) are used to pump inner (core), middle (shell), and outer (continuous phase) into the devices. Because of the use of biological sample, the syringe containing core fluid is usually cooled with an ice bag wrapped around the syringe. Once the device is filled with liquid, flow rates are adjusted for obtaining stable double emulsion drop formation. After drops exited the device, they continued to travel to a UV-crosslinking section where a 365 nm UV lamp (UVP Multiple-Ray Lamp, Fisher Scientific, Hampton, NH) crosslinked the polymer in the shell phase, producing microcapsules. Glass vials were typically used for capsule collection by inserting the end of the exit tubing into the vial prefilled with 10ml of PBS solution. Emulsion drops are typically collected for a period of 15-20 mins, depending on the need. Then the continuous fluid was replaced by using filtration tool to remove the original continuous fluid and resuspend the capsules in fresh solution containing 10 wt% glycerol and 2 wt% poly vinyl alcohol for osmotic balancing with the core fluid.

Drop production was visualized on a microscope equipped with a fast camera (Photron Mini AX100, Photron, San Diego, CA) capable of up to 540,000 frames/sec. Images of the double emulsion drops and capsules were analyzed with the freely available image analysis program, ImageJ (Rasband, W.S., U. S. National Institutes of Health, Bethesda, MD).

### FCS instrument

The FCS instrument was homebuilt using a multiple wavelength (637 nm, 561 nm, 488 nm, 405 nm) laser system (Stradus VersaLase 4, Vortran Laser); the laser is coupled through an optical fiber and used for excitation. A photodiode (PDA-155, Thorlabs) is used to monitor the laser power. An excitation DM (Di01-R405/488/561/635-25×36, Semrock) reflects excitation wavelengths and transmits emission wavelengths. We use a 60x water immersion objective (UPlanApo/IR, 1.20 NA, Olympus) to focus the excitation beam into a detection volume of ∼1.6 fL. The emission beam is directed through a 50 μm pinhole and split into red and green wavelengths by the emission DM (FF562-Di03-25×36, Semrock). The split beam then passes through bandpass filters (FF01-609/54-25 and FF02-525/40-25, Semrock) and is split again by 50/50 beamsplitters onto two avalanche photodiodes. We use four APD detectors (PD-050-CTD, Micro-Photon-Devices), two per color, for detection of photon counts. For this work, we used the 488 nm laser and the two APDs in the green emission channel.

The detection volume was determined by a calibration using solutions of Atto 488 dye at known concentrations. The concentration of a stock solution of Atto 488 was measured using a spectrometer (Nanodrop), and dilutions ranging from 500 pM to 100 nM were made from the stock then measured on FCS. Correlations were fitted to the 1-species model. The volume was calculated as the slope of a linear fit to the plot of inverse amplitude vs. concentration. The measured detection volume ranged from 1.5-1.8 fL. This variation could be due to slight differences in alignment and environmental factors (e.g. temperature) over time.

### FCS data acquisition and analysis

Using homebuilt software, arrival times of each photon were recorded on a counter card (National Instruments PCIe-6612). Correlations were calculated following the method outline in Reference ^31^. The cross correlation between the two green APDs was used as the autocorrelation function to avoid the afterpulsing effect.

Correlations were fitted to 1- or 2-species models to obtain amplitudes and diffusion times. For fits to correlation data with two species, we first fitted to the 1-species model to confirm the overall trend in diffusion time. Then the correlation data was fitted to the 2-species model with the two diffusion times τ_D1_ and τ_D2_ fixed at values indicated by the 1-species fit (the unbound and fully bound diffusion times). Typically, unbound τ_D1_ = 0.23 ms and bound τ_D2_ = 0.46 ms in homemade lysate and buffer; τ_D1_ = 0.35 ms and bound τ_D2_ = 0.76 ms in commercial lysate. Slight variations in measured diffusion times could be due to variations in detection volume/alignment. We noted earlier that the measured diffusion time of GFP is higher in the commercial lysate than in homemade lysate or buffer, perhaps due to interactions with components of the lysate or oligomerization in the buffer condition of the lysate. Thus, batch variations in the commercial lysate could also contribute to variations in measured τ_D1_ and τ_D2_.

## Author contributions

CL, EJL, and MJL performed FCS experiments. CL and EJL analyzed data. SAH produced cell-free lysates. CY and EJF performed encapsulation. CL, MJL, NNW, and TAL built the FCS instrument. NMH helped with tests of cell-free lysate and encapsulation. BDV contributed key reagents of cell-free lysates. CL, NNW, TAL, and MAC conceptualized the project. CL, SAH, CY, EJL, and TAL wrote the paper. All authors contributed intellectually to discussions of the results and reviewed and edited the paper.

## Acknowledgements

The authors would like to thank Sean Gilmore and Maxim Shusteff for initial planning of the project; Kansas Seung and Parker Moberg for helping build the sample stage; and Bonnee Rubinfeld for providing plasmids. This project was funded by Lawrence Livermore National Laboratory Laboratory-Directed Research and Development grant 21ERD039 and by the GUIDE program. The GUIDE program is executed by the Joint Program Executive Office for Chemical, Biological, Radiological, and Nuclear Defense (JPEO-CBRND) Joint Project Lead for Enabling Biotechnologies (JPL CBRND EB) on behalf of the Department of Defense’s Chemical and Biological Defense Program. This effort was in collaboration with the Defense Health Agency (DHA) COVID funding initiative. <u>Disclaimer</u>: The views expressed in this paper reflect the views of the authors and do not necessarily reflect the position of the Department of the Army, Department of Defense, nor the United States Government. References to non-federal entities do not constitute nor imply Department of Defense or Army endorsement of any company or organization. This work performed under the auspices of the U.S. Department of Energy by Lawrence Livermore National Laboratory under Contract DE-AC52-07NA27344.

---

**Supplementary Figure 1.**
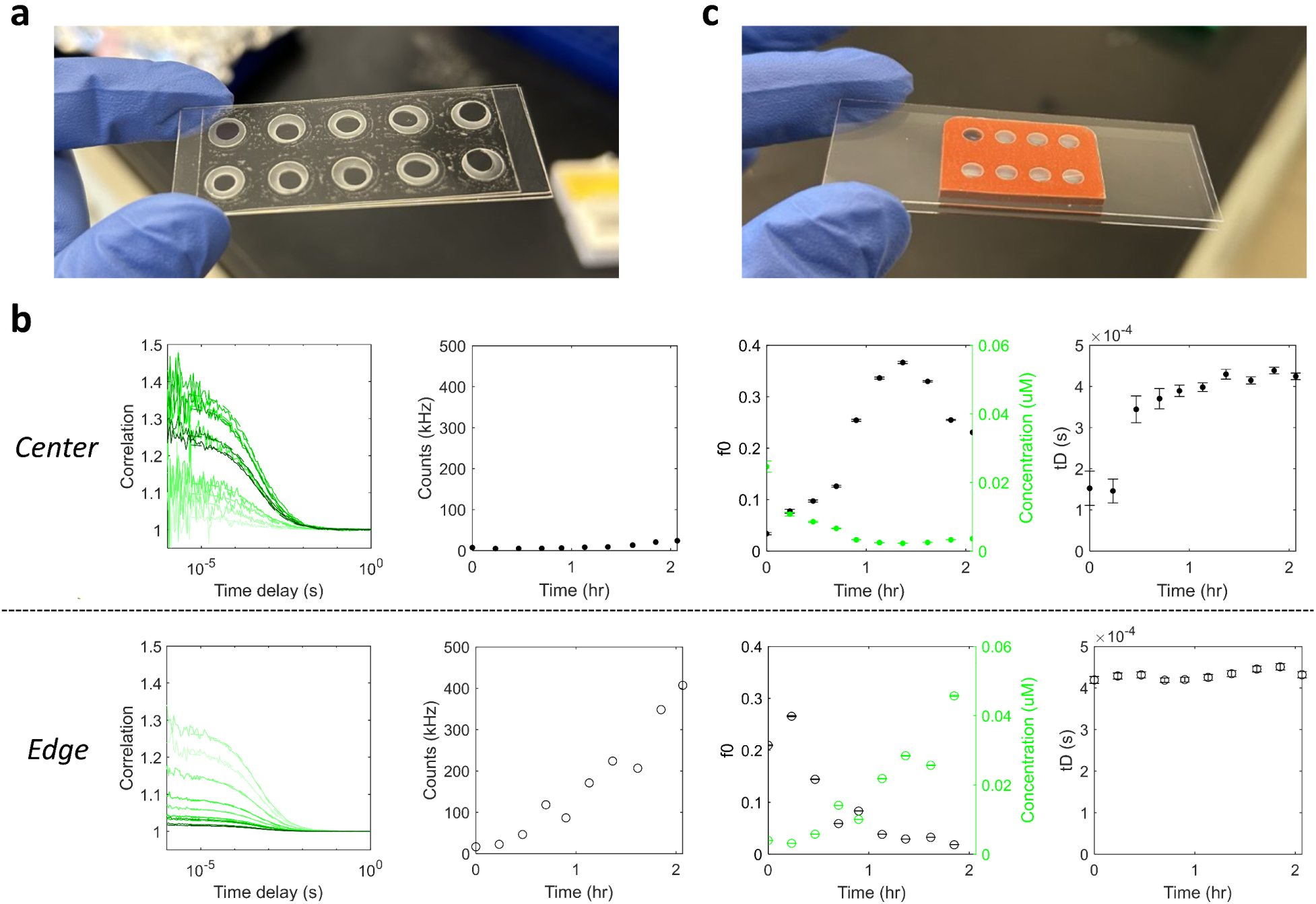
Sample chambers must provide sufficient oxygen for cell-free production of fluorescent proteins. **a.** A sample slide consisting of wells ∼120 um deep and 6.35 mm wide sealed between two coverslips. 2 µL of sample solution was deposited into each well, which formed a patch ∼3 mm in width (they appear as the dark circles inside each clear well) surrounded by residual air inside each well. **b.** FCS time series data acquired near the center (top) and edge (bottom) of the same well of a slide prepared as in (**a)**, demonstrating significantly faster and higher increase of GFP signal near the edge than near the center. 1 µg/ml GFP plasmid in commercial cell-free lysate was used for this data. Near the center, the GFP signal remained low for over 1 hr before rising slowly; correlation curves started out noisy with low amplitude since the signal was dominated by the lysate background; as GFP was slowly produced, the amplitude increased initially as the signal rose above lysate background, then decreased as the GFP concentration further increased; and the fitted diffusion times slowly increased from a value corresponding to the noisy lysate background to the value corresponding to GFP. In contrast, the GFP signal near the edge of the solution patch increased much faster, presumably because oxygen was more readily available near the border with air. **c.** A sample slide consisting of wells ∼1.8 mm deep and 3 mm wide between two coverslips. To provide uniform accessibility to air, we filled the bottom half of these wells with 6 µl cell-free reaction, leaving a layer of air above the solution. This configuration resulted in uniform GFP signal in all directions.

**Supplementary Figure 2.**
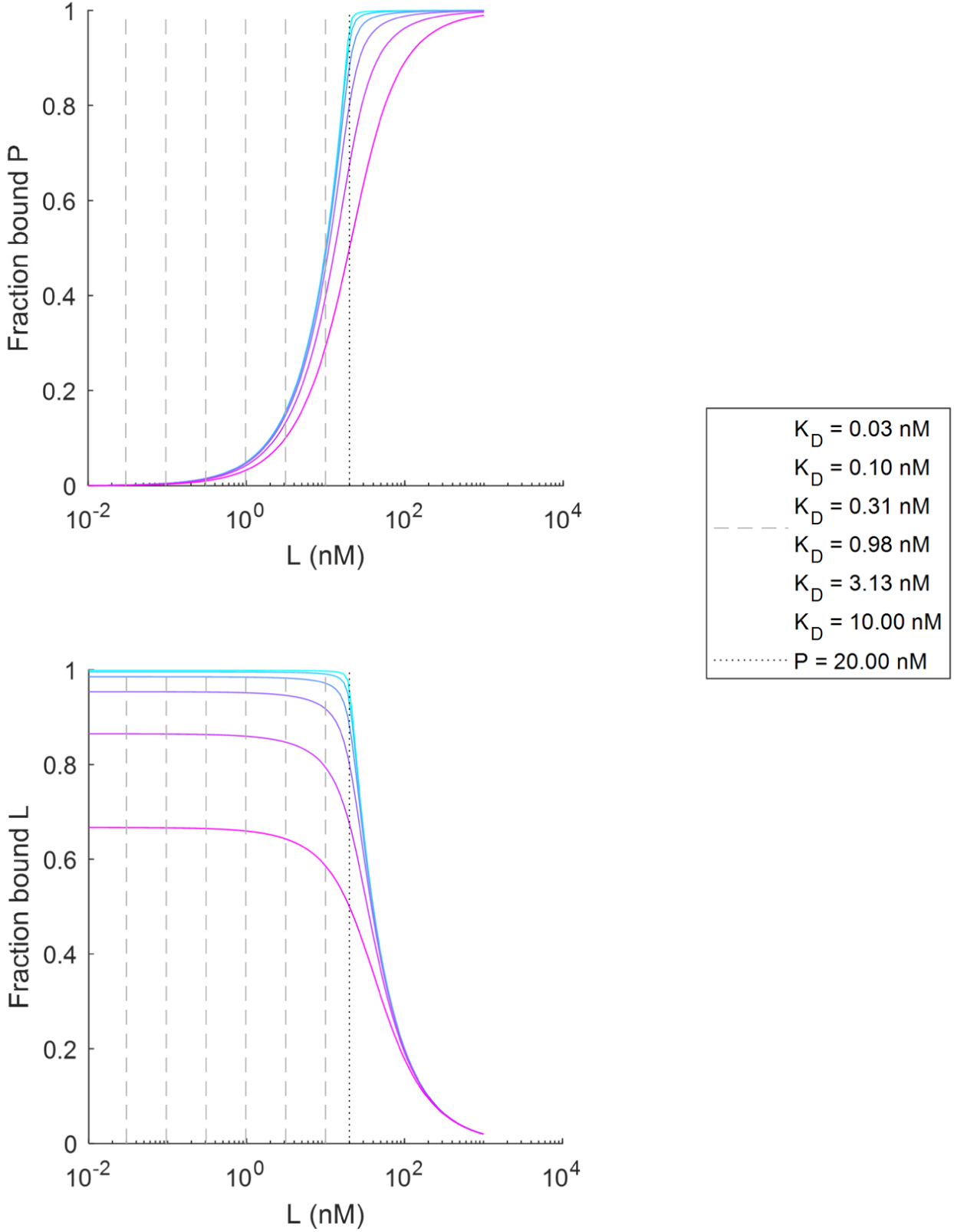
Accurate determination of *K*_D_ is not possible when the trace protein concentration is much higher than the true *K*_D_, as illustrated by simulated binding curves. In this example, the trace protein concentration is set to *P* = 20 nM, and binding curves corresponding to *K*_D_s ranging from 30 pM (cyan) to 10 nM (magenta) are simulated. Dashed lines represent the *K*_D_ values, and the dotted line represents the *P* value, for comparison. The top panel plots fraction of the trace protein bound as a function of ligand concentration (the canonical way of plotting a binding curve), and the bottom plots fraction of ligand bound as a function of ligand concentration. It is apparent that the binding curves corresponding to the lower picomolar *K*_D_s would be indistinguishable from each other in the presence of experimental noise, as expected since *P* = 20 nM is much higher than the *K*_D_s. In these cases, the measured *K*_D_ would be interpreted as the upper bound.

**Supplementary Figure 3.**
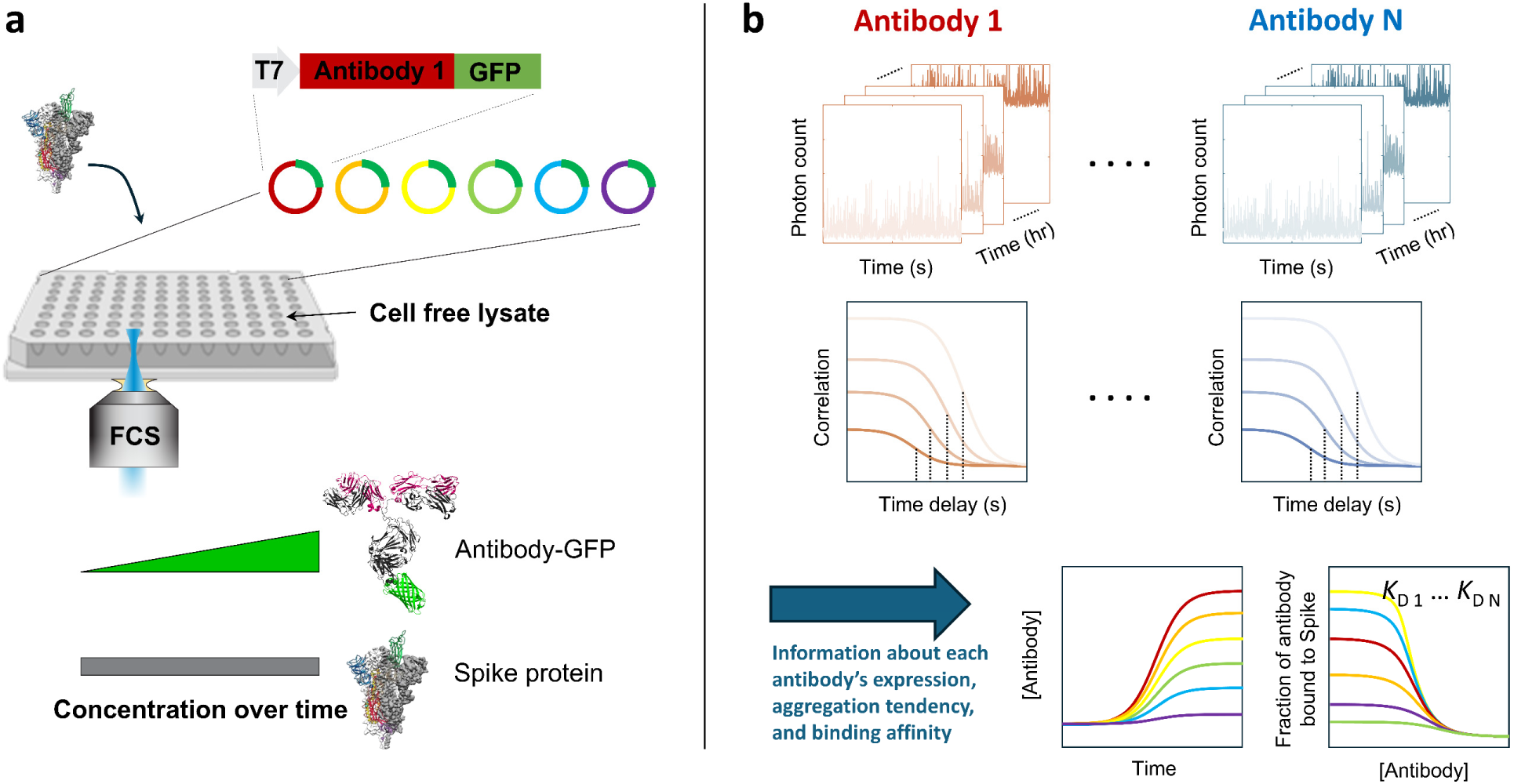
Approach for high throughput screening of candidate binders using the “one-pot reaction” method. **a.** In this illustration, different antibody or antibody fragment constructs are to be screened for binding to the COVID spike protein. GFP is a part of the antibody construct and provides the fluorescence signal for FCS. Each sample well contains the cell-free lysate, the purified spike protein, and a plasmid encoding one antibody-GFP candidate. FCS data is acquired in each well over cell-free expression time so that the concentrations of antibody-GFP increase while the concentration of the spike protein remains constant, analogous to the experiments presented in this paper in which GFP was expressed over time as the anti-GFP antibody remained constant. **b.** Overview of data analysis. Each candidate antibody would have a set of FCS data, e.g., 30 s of photon stream repeatedly acquired over 5 hr. Each 30 s segment of photon stream would yield a correlation curve from which the concentration of the antibody-GFP and the fraction bound are obtained. From here, each antibody would have an expression profile (concentration over time) and a binding curve.

## Notes

### Competing Interest Statement

The authors have declared no competing interest.

